# Cerebral oxygenation and cardiac output responses during short repeated-sprints exercise and modulatory effect of glucose ingestion

**DOI:** 10.1101/2022.12.05.519099

**Authors:** Paulo A. S. Armada-da-Silva, Hu Mingzhu, Wu Zongze, Wen Linjian, Feng Ruisen, Xinglin Zeng, Zhen Yuan, Zhaowei Kong

## Abstract

In this study we investigated changes in concentrations of cerebral oxyhaemoglobin (O_2_Hb), deoxyhaemoglobin (HHb), and total haemoglobin (tHb) during repeated sprints and their relationship with cardiac output. We also examined the effect of glucose ingestion and acute hyperglycaemia on cerebral haemoglobin responses. Ten young male participants ingested either a glucose drink (70 g) or a placebo before performing a set of 10 repeated 6-second sprints on a cycle ergometer. Heart rate, stroke volume, and cardiac output were measured continuously using impedance cardiography, while changes in O_2_Hb, HHb, and tHb in the frontal region of the cerebral cortex were measured using near-infrared spectroscopy (NIRS). The results showed that each sprint elicited a transient increase in O_2_Hb and to lesser extent HHb concentrations, which was enhanced with the number of sprint repetitions and correlated with cardiac output. After each sprint, O_2_Hb and HHb quickly returned to baseline, while cardiac output remained elevated. At the end of the repeated sprints, O_2_Hb was decreased to below pre-exercise levels, while HHb and tHb were elevated. After a recovery of 10-15 min, O_2_Hb returned to pre-exercise values in the placebo trial, but increased to above pre-exercise values (reactive hyperaemia) in the glucose trial. Our findings suggest that short sprint exercise increases O_2_Hb, HHb, and tHb levels during exertion in parallel with cardiac output. However, in addition to the transient increase in cerebral haemoglobin, a progressive decline in cerebral oxygen saturation occurs during repeated sprints. Glucose ingestion does not alter cerebral haemoglobin responses to sprint exercise but appears to be associated with faster recovery of O_2_Hb.

## Introduction

Studies investigating cerebral blood circulation have reported an increase in regional cerebral blood flow (rCBF) during dynamic exercise of moderate-to-severe exercise intensity [i.e., ∼ 70% maximal oxygen uptake (V O_2max_)]. However, when the work rate increases to maximal exhaustive levels, rCBF decreases towards baseline ^1,2^. In addition to augmented CBF, dynamic exercise also results in an increase in cerebral metabolism, as evidenced by a lowered cerebral metabolic ratio ^3,4^, indicating enhanced neural activity ^5^. This response is typically observed during exhaustive exercise ^6^, exercise in hyperthermic and hypoxic conditions, or exercise under increased mental effort ^4^.

In addition to changes in rCBF, dynamic exercise also affects cerebral oxygenation ^7^. Changes in concentration of oxyhaemoglobin (O_2_Hb) and deoxyhaemoglobin (HHb) in the cerebral tissue can be non-invasively measured with near-infrared spectroscopy (NIRS) with a reasonable spatial resolution (∼1-2 cm) and high time resolution. Two independent meta-analyses of NIRS studies have shown that O_2_Hb concentration in the frontal region of the cerebral cortex increases during submaximal exercise but decreases as exercise approaches maximal intensity, suggesting a mismatch between oxygen delivery and demand at the end stages of incremental maximal exercise ^8,9^. Such mismatch has been related with a cerebral vasoconstriction in response to exercise-induced hyperventilation and hypocapnia ^2,10^. The lowered cerebral oxygenation at near-maximal exercise, combined with muscle deoxygenation, may impair muscle recruitment and trigger fatigue ^2^.

Extremely short all-out sprint interval exercise (<10 s) is an attractive option for achieving cardiorespiratory fitness ^11^, but the cerebrovascular response to this type of exercise is unclear. One study showed that a 30-second all-out cycling exercise at normal ambient oxygen levels and in hypoxia resulted in a decrease in middle cerebral artery blood flow mean velocity (MCA*v*) and cerebral oxygenation, which were unrelated to changes in end-tidal carbon dioxide partial pressure (P_ET_CO_2_) ^12^. In addition, the MCA*v* decrease during the 30-second all-out cycling did not follow the changes in cardiac output and mean blood pressure, which increased during the exercise ^12^. In another study, participants performed seven 30-second exercise bouts on a cycle ergometer at a work rate of 150% maximal oxygen uptake, where O_2_Hb concentration increased during the first bout but decreased in the last three bouts, and HHb concentration increased from resting levels, without a clear correlation between cerebrovascular response and P_ET_CO_2_ ^13^. These two studies suggest that, unlike submaximal and maximal exercise, cerebrovascular response during supramaximal exercise is not strictly dependent on arterial CO_2_ levels, mean blood pressure, or cardiac output. However, the impact of repeated short sprint efforts on cerebral blood flow and oxygenation, and how it affects performance, remain unknown. It is worth noting that dynamic cerebral autoregulation, which is another major mechanism regulating cerebrovascular function, has a time delay similar to the duration of short sprint bouts (∼6 s) ^14^, which may impair its protective role in preventing a steep rise in cerebral blood flow and the risk of hyperperfusion during all-out exercise.

Blood glucose levels play a role in determining cerebral metabolism and CBF ^15^. Although glucose is the primary energy substrate utilized by the brain, acute hyperglycaemia has been shown to decrease CBF and the cerebral metabolic rate of both oxygen and glucose ^15,16^, although opposite findings have been reported ^17^. On the other hand, glucose ingestion has been shown to improve cerebral glucose uptake and reduce perceived exertion during prolonged exercise, suggesting a potential ergogenic effect of glucose through central mechanisms ^18^. Hence, we also tested the effect of acute glucose intake and subsequent rise in glycaemia in cerebral haemoglobin concentration changes during repeated sprints and how it relates with performance.

Therefore, this study aimed to investigate changes in cerebral O_2_Hb, HHb, and tHb during the performance of repeated cycling sprints and their relationship with both cardiac output and power output. The effect of acute hyperglycaemia on cerebral (prefrontal) oxygenation during the repeated sprints was also investigated. A NIRS probe scanning the entire prefrontal cortex was used to assess brain oximetry, which allows compensating for interindividual cerebral circulatory regional differences and to investigate differences in cerebrovascular responses to short intermittent cycling sprints across the frontal region of the cerebral cortex.

## Methods

### Participants and experimental design

Eleven healthy male adults volunteered (Table 1) to participate in this study after providing written informed consent. Participants were free of any metabolic, cardiovascular, or neurological disease and did not regularly participate in exercise training. The study protocol was reviewed and approved by the ethics review board of the University of Macao (Special Administrative Region of Macao, China) and conformed to the ethical principles for research involving human participants (approved number: BSERE22-APP011-FED).

**Table 1:**
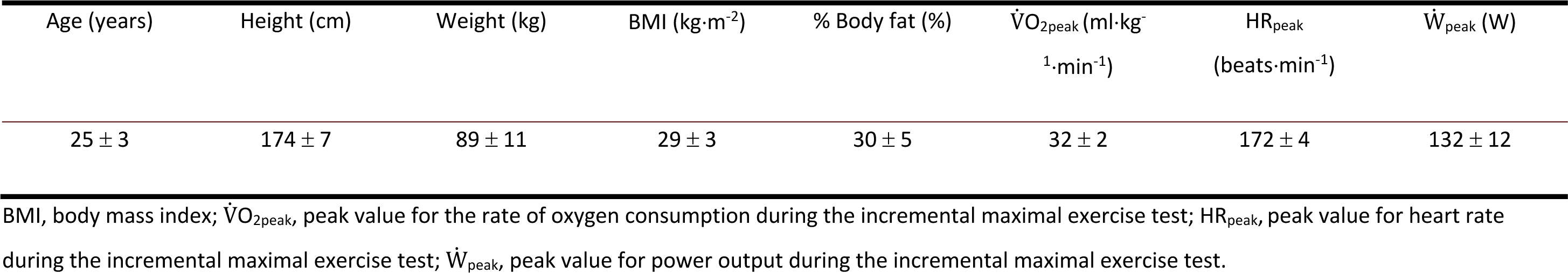
Participants characteristics. Data are presented as mean ± standard deviation; n = 11.

| Age (years) | Height (cm) | Weight (kg) | BMI (kg·m<sup>-2</sup>) | % Body fat (%) | $\dot{V}O_{2peak}$ (ml·kg<sup>-1</sup>·min<sup>-1</sup>) | HR<sub>peak</sub> (beats·min<sup>-1</sup>) | $\dot{W}_{peak}$ (W) |
| --- | --- | --- | --- | --- | --- | --- | --- |
| 25 ± 3 | 174 ± 7 | 89 ± 11 | 29 ± 3 | 30 ± 5 | 32 ± 2 | 172 ± 4 | 132 ± 12 |

BMI, body mass index; $\dot{V}O_{2peak}$ , peak value for the rate of oxygen consumption during the incremental maximal exercise test; HR<sub>peak</sub>, peak value for heart rate during the incremental maximal exercise test; $\dot{W}_{peak}$ , peak value for power output during the incremental maximal exercise test.

Each participant visited the laboratory on three separate occasions at least one week apart. In the first visit, each participant performed a maximal incremental cycling exercise to determine the individual VO_2max_. In the following visits, each participant performed a repeated sprint exercise test on a cycle ergometer after ingesting 70 g of glucose (GLU) or a taste-matched non-caloric placebo (PLA), in a randomised counterbalanced and single-blind design.

### Instrumentation and measures

### Maximal incremental test

The maximal incremental test was carried out on a cycle ergometer (Monark 839E, Varberg, Sweden). After a warm-up at 25 W for 3 min, participants started cycling at 50 W and a cadence of 60 ± 5 revolutions per minute (rpm), with the workload increasing by 25 W every 3 min. The test was completed when the participant reached volitional exhaustion, with a respiratory exchange ratio (RER) ≥ 1.15, ratings of perceived exertion (RPE) ≥18, and peak HR ≥ 90% of age-predicted maximum (220-age) ^19^. Respiratory gases were collected by the Vmax Encore System (CareFusion Corp., San Diego, CA, USA) throughout the test, and V O_2max_ was calculated as the highest oxygen consumption value averaged over 15 s before the end of the test ^11^.

### Repeated sprints cycling exercise

During the two experimental sessions, participants performed a repeated sprint set of 10 repetitions of 6 s maximal sprints interspersed by 24 s resting, performed on a cycle ergometer (Monark Ergomedic 894E, Monark Exercise AB, Vansbro, Sweden). Each participant sat upright on the cycle ergometer keeping the head as steady as possible with the help of a neck cushion to minimise motion artefacts during NIRS recordings. Before the repeated sprints, a 5-min warm-up, cycling at a power output of 60 W (resistance, 1 kg; cadence, 60 rpm) was completed. During the sprints, participants were instructed to use the same ready position, sitting upright and with one leg ready to push powerfully against the pedal and maximally accelerate the cycling crank at the “go” signal. Each participant was free to choose his leading leg in the first session and was asked to use the same leg to initiate all subsequent sprint repetitions. Five seconds into the next sprint, participants adopted the ready position and awaited the verbal go command at the end of a 3-s countdown. A weight equal to 6.5% body mass was applied to the flywheel controlled by the Monark Anaerobic Test Software at the instant the pedalling cadence reached 90 rpm. The controlling software recorded the mean power output for each sprint.

### NIRS

A portable 2 x 4-channel continuous-wave NIRS system (Octamon, Artinis Medical Systems, Einsteinweg, The Netherlands) was used to measure changes (Δ) in O_2_Hb, HHb, and tHb concentrations using two wavelengths at 750 and 850 nm. Data were acquired with the OxySoft software (OxySoft, Artinis Medical Systems, Einsteinweg, The Netherlands) at a frequency of 50 Hz with a differential path length factor of six. The optodes were attached to a dark ambient light-protecting probe holder, tightly secured to the head by adjustable headbands and placed over the forehead and the underlying prefrontal cortex (four channels on each side). The two light detectors were aligned on Fp1 and Fp2 locations of the 10-20 system for the electroencephalography and surrounded by four light-emitting optodes with an inter-optode distance of 35 mm, providing a total of eight NIRS channels.

### Impedance cardiography

Cardiac output was measured continuously during the sessions by signal morphology impedance cardiography (PhysioFlow PF-05, Manatec biomedical, Paris, France). After skin preparation to minimise electrical resistance, six Ag/AgCl electrodes (Hewlett Packard 40493 E) were placed on the chest wall along the xiphoid process and at the V1 and V6 precordial locations, and at the base of the neck to record the thoracic impedance and electrocardiographic signals. Resting blood pressure was measured by automatic oscillometry to calibrate the cardiac impedance data acquisition following the manufacturer instructions. Stroke volume and cardiac output were recorded for each cardiac cycle and measured continuously throughout the trial.

### Blood glucose and lactate

Blood glucose measurements were collected with a glucometer (ACCU−CHEK Performa, USA) and the proper test strips (F. Hoffmann-La Roche AG, Basel, Switzerland). Blood lactate was also measured with the Lactate Pro 2 handheld device and respective test strips (AKRAY Europe B.V., Amstelveen, the Netherlands). Both blood analytes were measured in duplicate from capillary blood drawn from a lanceted fingertip at three time points: 1) at baseline, soon after arrival to the laboratory; 2) immediately before starting the exercise protocol; and 3) immediately after completion of the repeated sprints exercise. The duplicate blood sample measurements were averaged for analysis.

### Procedures

Participants arrived at the laboratory between 8:00 and 9:00 am after a 12-hours fast. Soon after arrival, participants ingested 300 ml of a drink containing 70 g of glucose (Cambridge Isotope Laboratories, Cambridge, MA, USA) or a taste-matched placebo containing artificial sweeteners (Truvia Calorie-Free Sweetener; Cargill, IL, USA). Next, they rested sitting for one hour. Fifteen minutes to the end of the one-hour resting, participants were fitted with the cardiac output, and NIRS instrumentations and the two devices were set up for acquisition. The trial started with 10 minutes of seated resting, while wearing a cushion around the neck to reduce head movement.

After the 10-min rest, participants mounted the cycle ergometer and completed the 5-min warm-up followed by the 10 x 6-s repeated sprints. After exercise, there were another 10 minutes resting. Ratings of perceived exertion were obtained at the end of the fifth and tenth sprint repetitions using the Borg’s scale.

### Data analysis

The NIRS data pre-processing was performed using the Homer2 toolbox ^20^ for Matlab (Matlab version 9.5.0, release R2018b, The Mathworks Inc., Natick, Massachusetts, USA). First, light absorption data were converted into optical density and haemoglobin concentrations using the modified Beer-Lambert law with a differential path length factor of six. Next, all NIRS signals were low pass filtered at a cut-off frequency of 0.5 Hz to remove heartbeat-associated changes in O_2_Hb, HHb and tHb, using a finite impulse response fourth-order Butterworth filter, implemented in the forward and inverse directions to avoid phase filtering artefacts ^20^. After filtering, the whole NIRS signal was epoched into the individual sprints, starting at minus 5 s and ending at 25 s after the sprint start. The median was used to quantify the changes in haemoglobin concentrations in the time intervals 0-15 s and 15-25 s, to measure the response during the sprints and during the rest interval between sprints, respectively. The median was chosen because of its higher robustness against potential biasing effects caused by motion artefacts and outliers. The time interval 0-15 s was selected following a visual inspection of the transient ΔO_2_Hb response accompanying the performance of the sprints (Fig. 3). Due to the non-normal distribution of some ΔO_2_Hb, ΔHHb, and ΔtHb values (Supporting information, Figs. S1-S3 and Table S1), a natural log-transformation was applied to these variables for the purpose of statistical analysis.

### Statistical analysis

The Shapiro-Wilk test was used to verify the normal distribution of the data. To assess the responses to repeated sprint exercise, a repeated measures ANOVA with a complete factorial design (cerebral oximetry data: 2 trials x 8 channels x 10 sprint repetitions; power output and cardiovascular data: 2 trials x 10 sprint repetitions) was conducted. The Greenhouse-Geiser correction was applied to within-subject factors violating the sphericity assumption (Mauchly’s test p-value < 0.05). Post hoc tests were performed employing one-way ANOVA with Bonferroni’s correction for family-wise error rate. Planned contrasts were performed using polynomial fitting. This approach treated the repeated sprints as a quantitative variable with even spacing. While the polynomial contrast analysis does not directly compare two specific sprint repetitions, it provides valuable information about the overall trend effect of the repeated sprints on the response variables. In case of a significant linear contrast, backward differences contrasts were computed (the contrast between two consecutive sprint repetitions, including resting). Further contrasts between the first repeated sprint and the following sprints were also calculated. One sample *t*-test was used for testing the significance of post-exercise mean ΔO_2_Hb, ΔHHb and ΔtHb against pre-exercise values (since continuous NIRS measurements are relative, O_2_Hb and HHb baseline concentrations are set to zero). Correlation values were computed using the Pearson’s coefficient. All statistical tests were performed in RStudio (v. 1.3.1093, 2009-2020 RStudio, PBC) using functions from Rstatix (v. 0.7.0) and R base (v. 4.0.3) packages. Unless otherwise stated, all data are reported as mean ± standard deviation (SD). Significance was accepted at p < 0.05.

## Results

Due to large noise affecting the NIRS signals, one participant was excluded from the final analysis, resulting in a final sample size of 10 participants. In another participant, the NIRS signals pertaining to three of the sprints in the two trials were removed from analysis because of significant motion artefacts. The missing data (48 out of 1600 data points) were imputed using the average values from the same individual, channel, and the other sprints. One participant in the PLA trial had missing mean power output data for the two final repeated sprints due to equipment failure. The last recorded mean power output was used to replace the missing values.

### Power output and RPE

Figure 1 displays the mean power output for each sprint and trial. The mean power output was found to be higher during the GLU trial than the PLA trial [GLU: 648 ± 183 W; PLA: 558 ± 161 W; F_(1, 9)_ = 6.312, p = 0.033, η^2^ = 0.098], particularly in the later sprint repetitions [trial x sprint repetition interaction effect: F_(2.79, 25.12)_ = 12.314, p < 0.000, η^2^ = 0.203]. According to one-way ANOVA, mean power output was significantly different between GLU and PLA in the first (PLA higher than GLU), seventh and tenth sprint repetitions (Bonferroni’s corrected p < 0.05, Fig. 1). At the end of the fifth repetition, perceived exertion was moderate (GLU: 11.9 ± 2.1; PLA: 13.2 ± 1.9 units), but RPE reached near-maximal scores at the end of the sprint set (GLU: 19.3 ± 2.1; PLA: 19.3 ± 0.5 units; main effect of sprint repetition, F_(1, 9)_ = 232.66, p < 0.000, η^2^ = 0.856), with no differences between trials.

**Figure 1:**
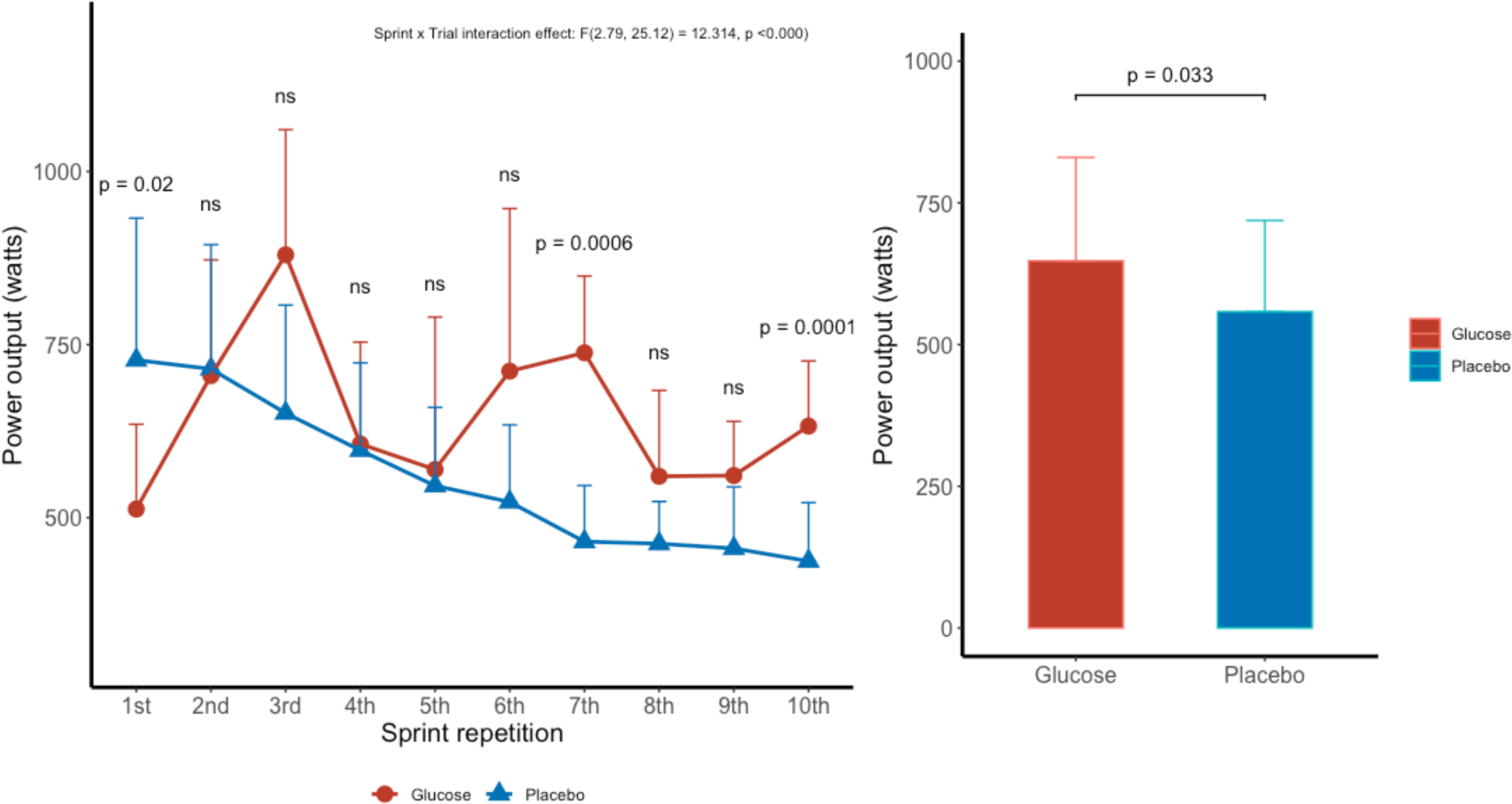
Mean power output during each repeated sprint for GLU (red line) and PLA (blue line) trials (left panel). Pooled mean power output for GLU (red bar) and PLA (blue bar) trials (right panel). Data presented as mean ± SD, ns indicates non-significant difference, sample size: n = 10.

### Blood glucose and lactate

At baseline, blood glucose concentration was similar in the two trials (GLU: 5.7 ± 0.5 mmol·l^-1^; PLA: 5.9 ± 0.6 mmol·l^-1^; p = 0.310). However, following glucose ingestion, the blood glucose concentration rose and was significantly higher in the GLU trial compared to the PLA trial prior to exercise (GLU: 8.1 ± 1.7 mmol·l^-1^; PLA: 6.1 ± 0.8 mmol·L^-1^; p = 0.016). By the end of the repeated sprints, the blood glucose concentration returned to similar levels in both trials (GLU: 6.5 ± 1.6 mmol·l^-1^; PLA: 5.9 ± 0.9 mmol·l^-1^; p = 0.254).

At baseline, the blood lactate concentration was at the same level in the GLU and PLA trials (GLU: 1.7 ± 0.9 mmol·l^-1^; PLA: 1.4 ± 0.4 mmol·l^-1^; p = 0.335). However, after glucose ingestion, the blood lactate concentration increased significantly (GLU: 2.4 ± 0.8 mmol·l^-1^; PLA: 1.6 ± 0.5 mmol·l^-1^; p = 0.023). Immediately after the repeated sprints, the blood lactate concentration was markedly elevated and significantly higher in the PLA trial than in the GLU trial (GLU: 10.8 ± 1.1 mmol·l^-1^; PLA: 11.7 ± 1.4 mmol·l^-1^; p = 0.047).

### Heart rate, stroke volume, and cardiac output

Figure 2 shows the changes in heart rate, stroke volume and cardiac output during the repeated sprints. The heart rate increased progressively throughout the sprints set, reaching its highest value of 163 ± 5 beats·min^-1^ in the final sprint [main effect for sprint repetition, excluding resting values: F_(3.45, 31.04)_ = 347.947, p < 0.000, η^2^ = 0.884], with no significant difference between GLU and PLA [F_(1, 9)_ = 0.018, p = 0.897, η^2^ < 0.000].

**Figure 2:**
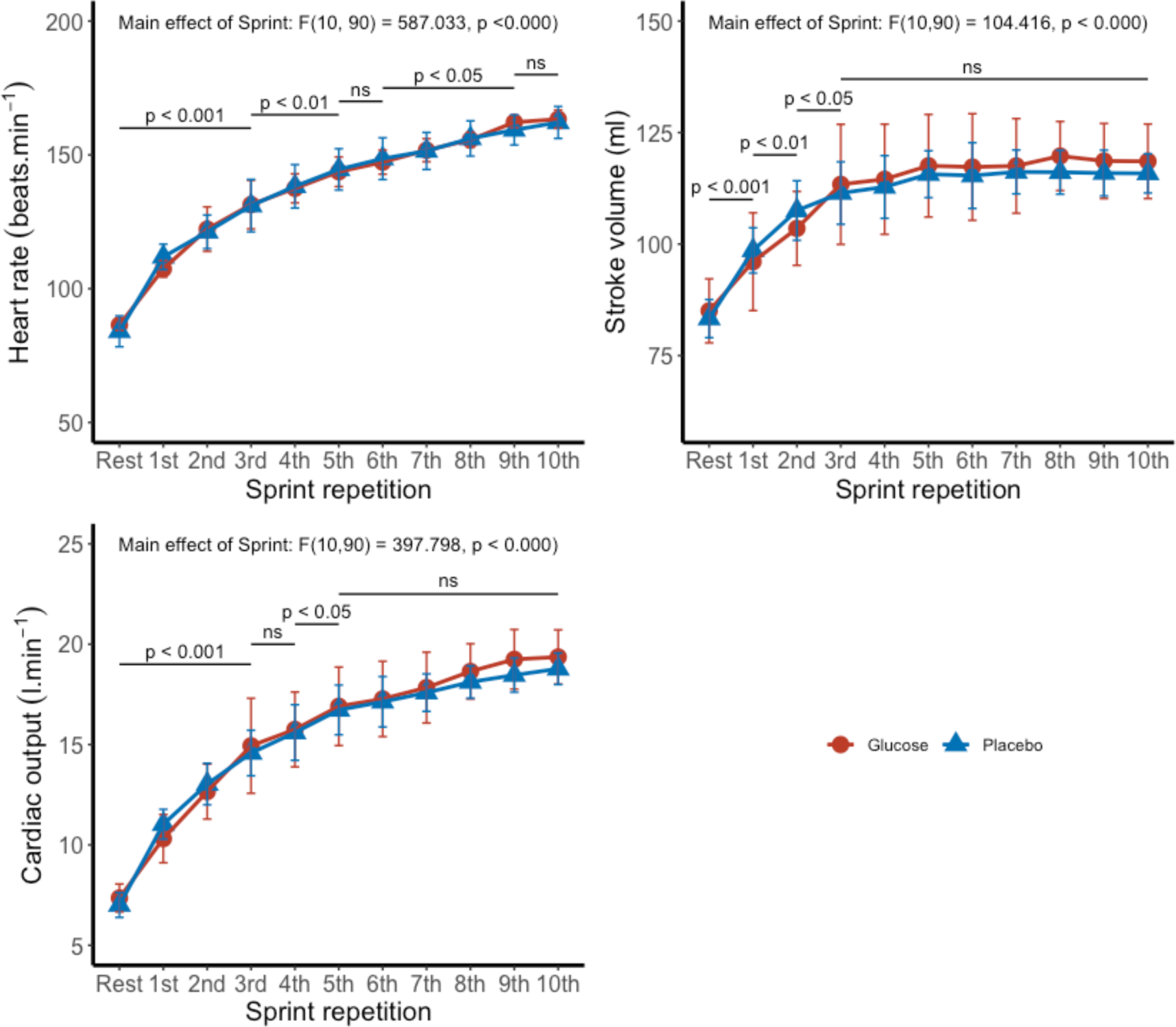
Heart rate, stroke volume and cardiac output (mean ± SD) at rest and during each repeated sprint for GLU (red lines) and PLA (blue lines) trials. The insert is the output of the two-way repeated measures ANOVA for the significant main effect of sprint repetition (excluding the resting values). P-values are from backward difference contrasts, comparing each sprint repetition with the immediately preceding one, including the first sprint and resting values. Sample size: n = 10.

The stroke volume increased from 84.2 ± 5.8 ml at rest to 97.3 ± 8.4 ml during the first sprint, but remained at a plateau after the fourth repetition [main effect for sprint repetition, excluding resting values: F_(2.47, 22.25)_ = 39.546, p < 0.000, η^2^ = 0.380]. There was no significant difference in stroke volume between GLU and PLA trials [F_(1, 9)_ = 0.195, p = 0.669, η^2^ = 0.005].

Cardiac output increased in a curvilinear pattern during the repeated sprints, rising from a resting value of 7.2 ± 0.7 l·min^-1^ to a cardiac output of 10.7 ± 1.0 l·min^-1^ in the first sprint and 19.1 ± 1.1 l·min^-1^ at the end of the repeated sprints [main effect for sprint repetition, excluding resting values: F_(2.59, 23.31)_ = 214.561, p < 0.000, η^2^ = 0.792]. There was no significant difference in cardiac output between GLU and PLA [F_(1, 9)_ = 0.280, p = 0.610, η^2^ = 0.005], as well as no interaction effect between trial and sprint repetition [F_(1.93, 17.38)_ = 1.122, p < 0.346, η^2^ = 0.025].

During the time interval between sprints, cardiac output remained high, but decreased slightly, although significantly, from the level attained during the sprints [overall mean ± SD during sprinting, 16.2 ± 2.8 l·min^-1^; in between sprints, 16.1 ± 2.8 l·min^-1^; F_(1, 9)_ = 6.084, p = 0.036, η^2^ = 0.003]. Polynomial contrasts showed a significant linear and quadratic effect of sprint repetition on heart rate, stroke volume, and cardiac output responses. Several backward differences contrasts were also significant (see Fig. 2 for details).

### Brain oximetry

Table 2 presents the results of ΔO_2_Hb, ΔHHb, ΔtHb during the repeated sprints and the rest periods between the sprint repetitions.

**Table 2:**
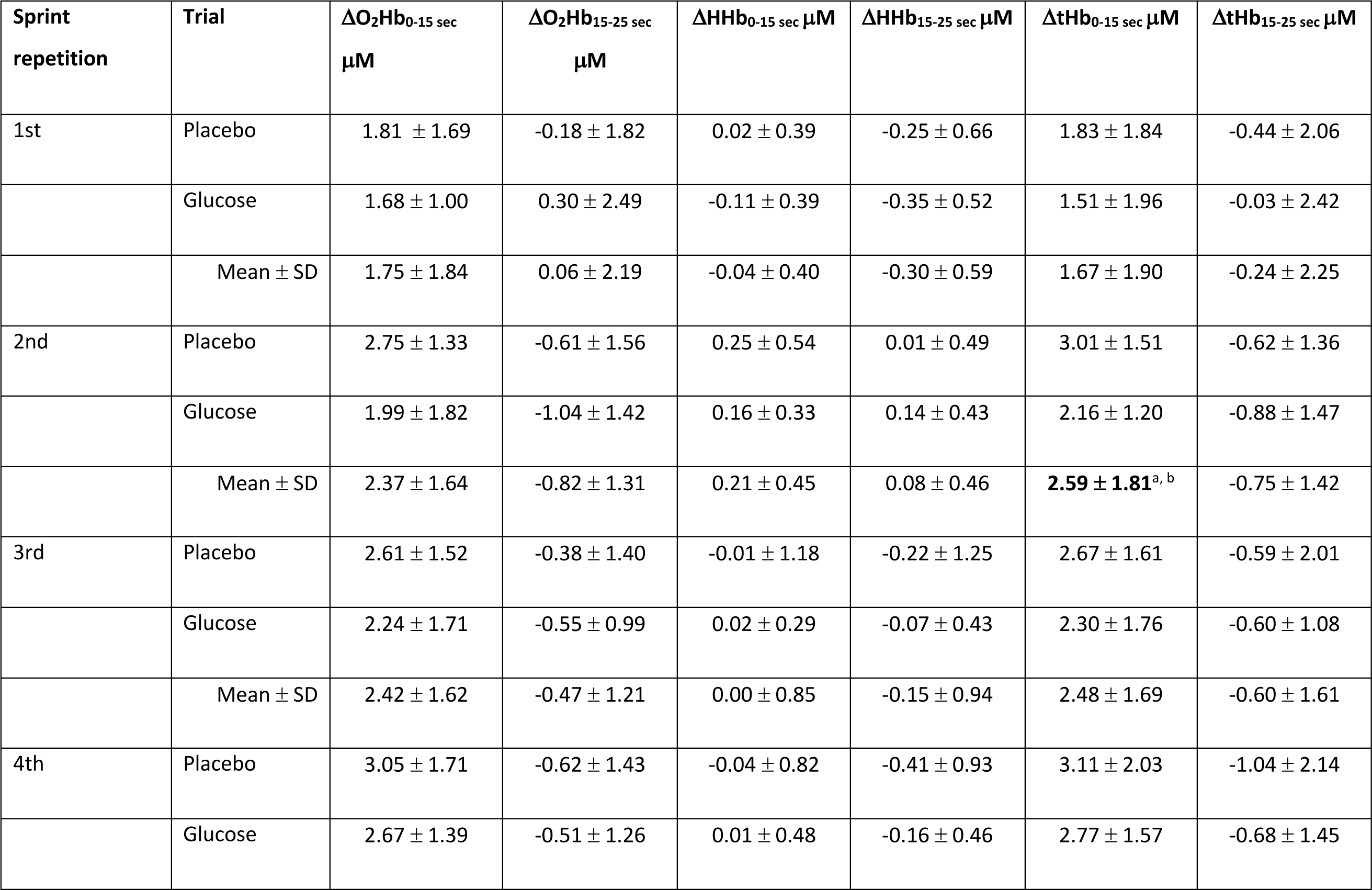
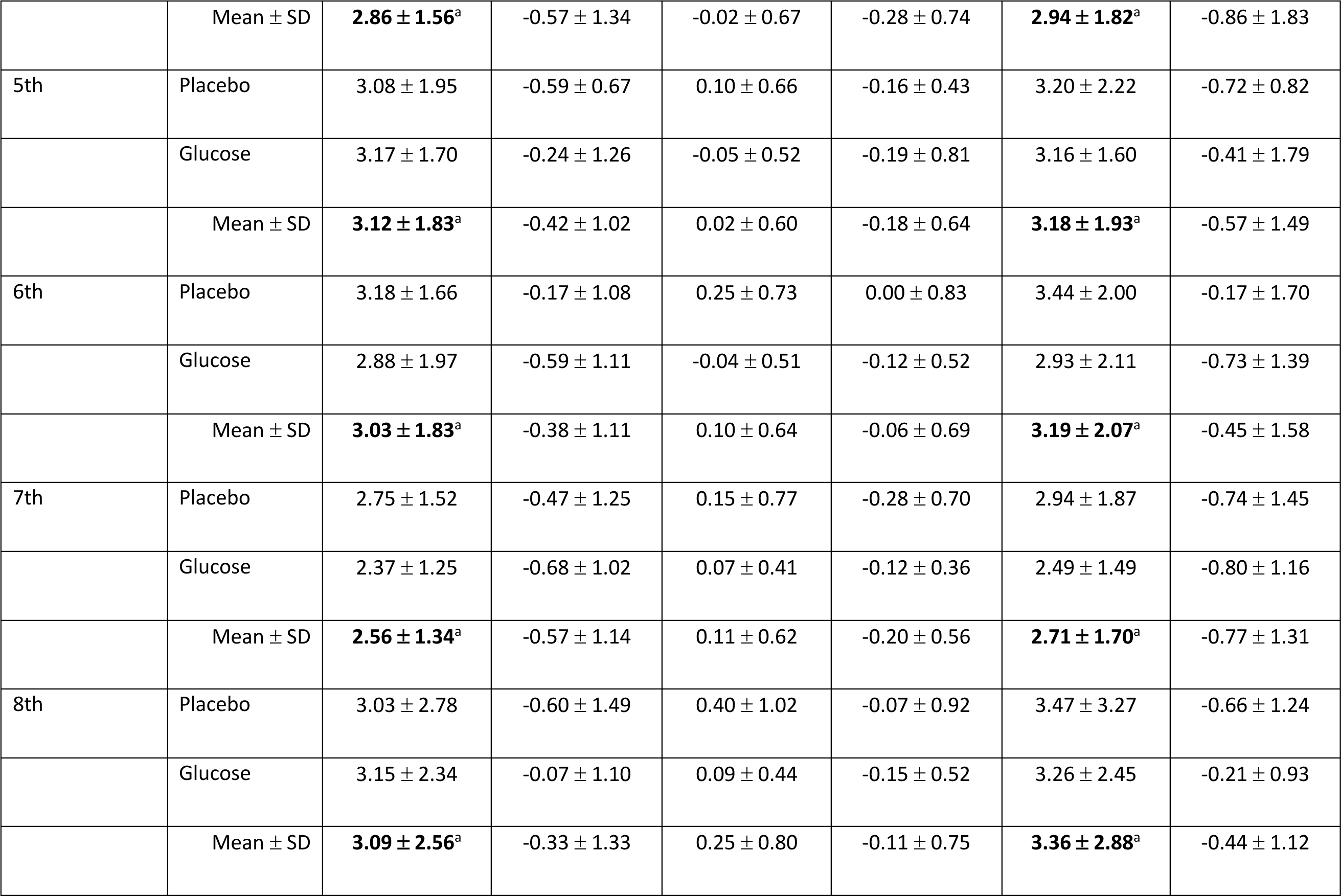
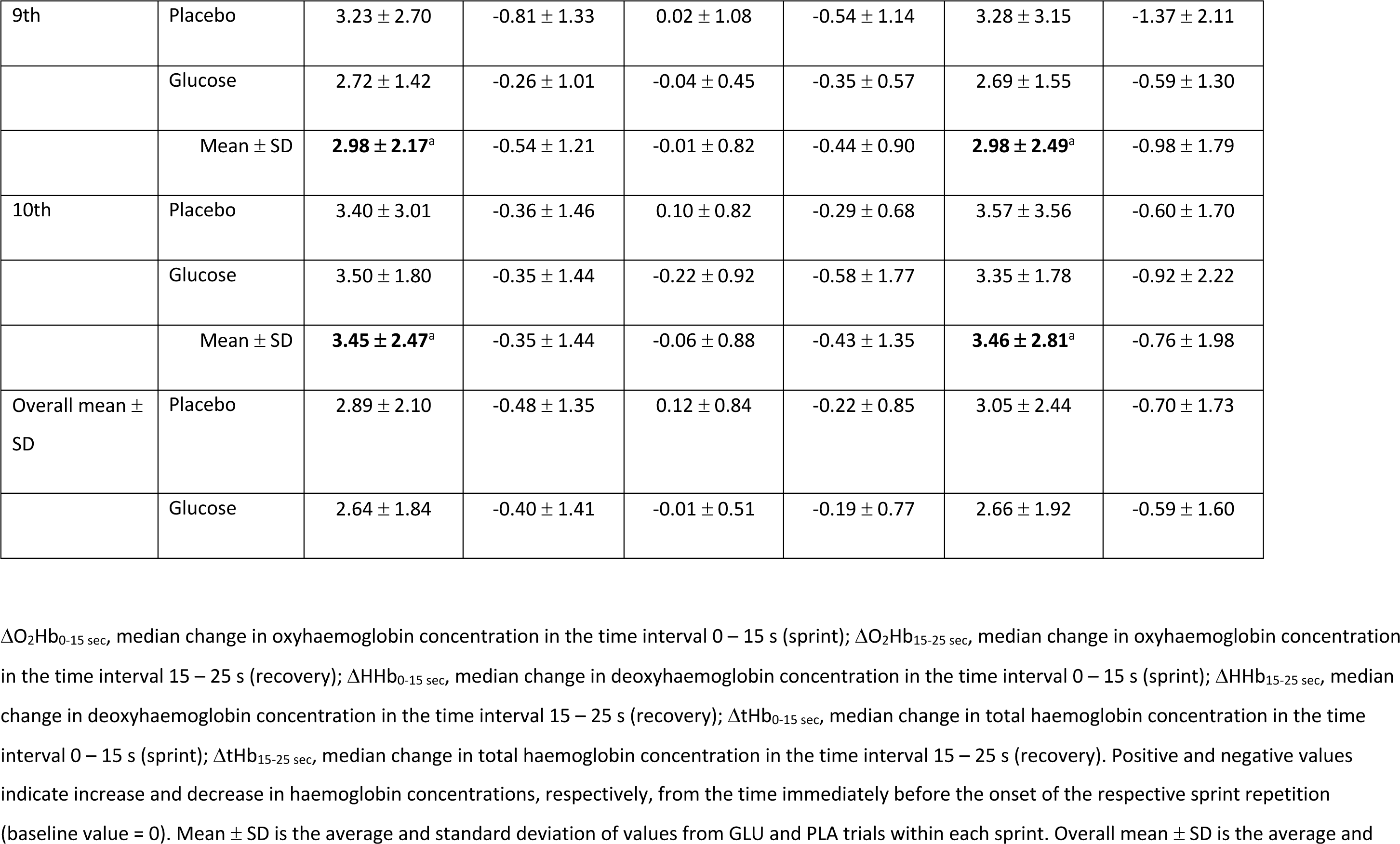
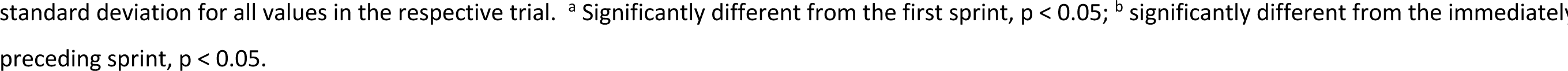
Mean ± SD values for oxyhaemoglobin, deoxyhaemoglobin, and total haemoglobin concentration changes during the repeated sprints; n = 10.

### Oxyhaemoglobin

During the sprints, O_2_Hb levels rapidly increased, reaching its peak at the end of each sprint (Fig. 3; Supporting information, Fig. S4). The O_2_Hb response to the sprints increased with the increasing number of sprint repetitions [main effect of sprint repetition: F_(9, 81)_ = 3.380, p = 0.001, η^2^ = 0.074]. However, there was no effect of glucose ingestion [F_(1, 9)_ = 0.930, p = 0.360, η^2^ = 0.005] or NIRS channel [F_(7, 63)_ = 10.222, p = 0.979, η^2^ = 0.002] on this response. Polynomial fitting demonstrated a significant linear effect of sprint repetition on ΔO_2_Hb, which did not improve with higher-order components. Additionally, contrast analysis showed significantly lower ΔO_2_Hb in the first sprint compared to the fourth and subsequent sprints (p < 0.05). After each sprint, O_2_Hb concentrations returned to pre-sprint levels or lower (Fig. 3, Table 2). There were no significant differences in ΔO_2_Hb values between trials, channel, or sprint repetitions for the rest periods between the sprints.

**Figure 3:**
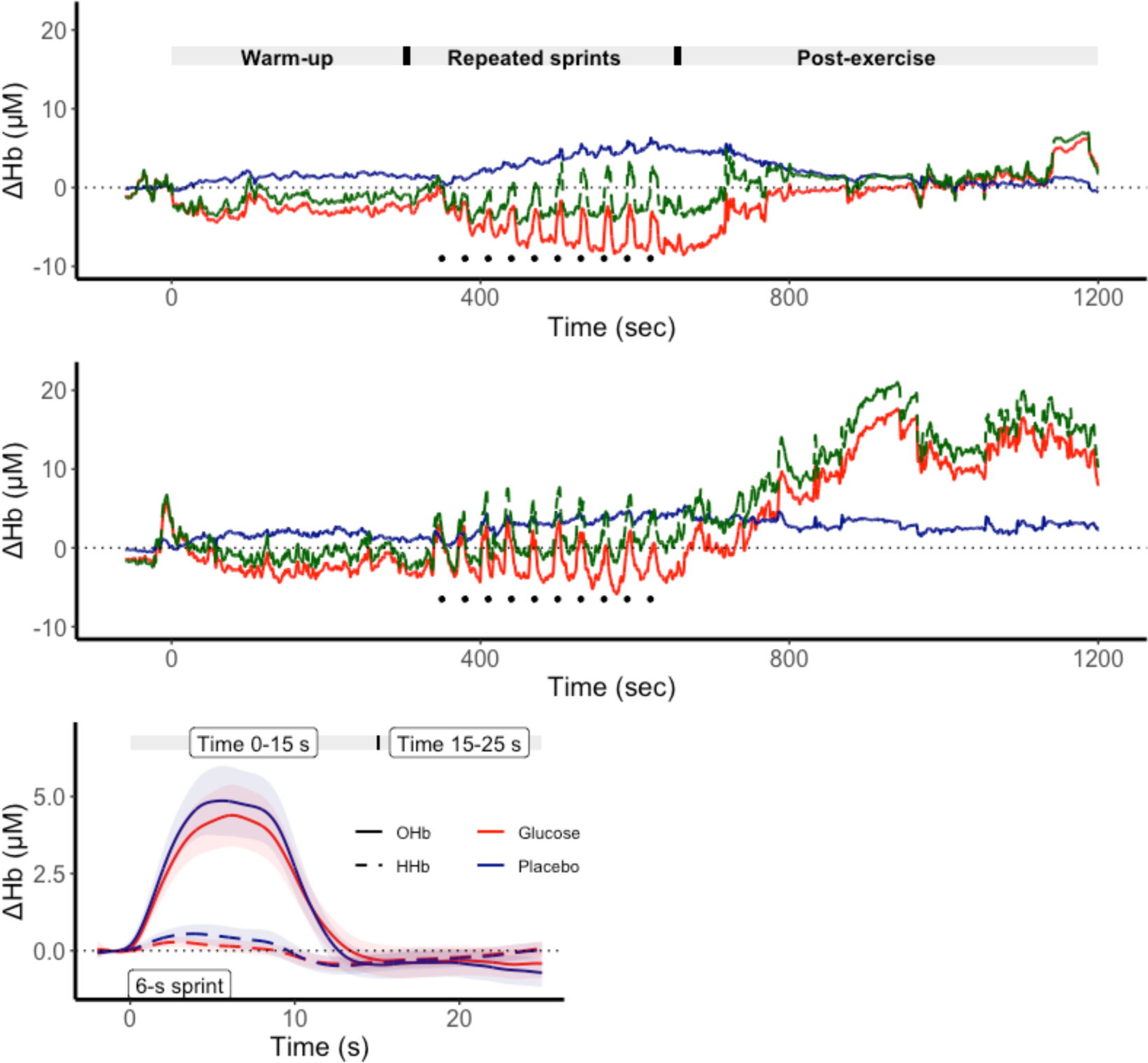
Changes in O_2_Hb, HHb and tHb during the repeated sprints exercise. Top and middle graphs display the concentration changes of O_2_Hb (red line), HHb (blue line) and tHb (green line) recorded in two participants. The records start immediately before the warm-up. In both participants, a transient increase in O_2_Hb, HHb, and tHb during each sprint (black dots) is clearly observed. In the top graph, there was a notable decrease in O_2_Hb concentration and increase in HHb throughout the repeated sprints exercise, followed by a return of both O_2_Hb and HHb towards pre-exercise values during the recovery period. The participant in the middle graph exhibited lesser changes in O_2_Hb and HHb concentration during exercise but displayed substantial post-exercise cerebral hyperaemia. The bottom panel presents the overall average for ΔO_2_Hb and ΔHHb during the repeated sprints, revealing the rapid increase in cerebral haemoglobin concentration during the sprint and its quick decrease soon after the end of the sprint. Each tracing is the mean of 10 participants, 10 sprints and 8 channels. The shadowed areas represent the standard error of the mean; sample size: n = 10.

### Deoxyhaemoglobin

In comparison to the changes in O_2_Hb, the changes in HHb in response to sprinting were less pronounced and not affected by the sprint repetitions [F_(9,81)_= 1.446, p = 0.182, η^2^ = 0.01] (Table 2, Fig. 3; Supporting information Fig. S4). Also, there were no significant differences in the changes of HHb between the two trials [F_(1, 9)_= 0.476, p = 0.508, η^2^ = 0.000] (Table 2) or between the NIRS channels [F_(7,63)_= 1.531, p = 0.173, η^2^ = 0.02]. During the rest interval between the sprints, HHb concentration returned to pre-exercise levels with no differences observed between the trials [F_(1, 9)_= 0.287, p = 0.605, η^2^ = 0.000] or NIRS channels [F_(7,63)_= 1.162, p = 0.337, η^2^ = 0.01], but with a significant main effect of sprint repetition [F_(9, 81)_= 2.152, p = 0.034, η^2^ = 0.012]. Polynomial contrasts revealed only marginal linear (p = 0.052) and quadratic (p = 0.057) effects of the repeated sprints in ΔHHb in the rest interval between the sprints.

### Total haemoglobin

The changes in tHb followed a similar pattern to that observed for O_2_Hb, with a marked increase during each sprint, followed by a rapid decline towards the baseline immediately after the sprint. Glucose ingestion [F_(1, 9)_= 0.970, p = 0.350, η^2^ = 0.007] and NIRS channels [F_(7, 63)_= 0.403, p = 0.897, η^2^ = 0.006] had no effect on the changes in tHb. A significant difference was observed in ΔtHb across the repeated sprints [F_(9, 81)_= 2.796, p = 0.007, η^2^ = 0.054], with a significant linear effect (p = 0.008) and lower ΔtHb observed during the first sprint compared to most of the subsequent sprints (0.003 < p < 0.046). No differences in ΔtHb during the time interval between sprints were observed between trials [F_(9, 81)_= 2.017, p = 0.189, η^2^ = 0.002], NIRS channels [F_(7, 63)_= 1.016, p = 0.429, η^2^ = 0.017], or sprint repetitions [F_(9, 81)_ = 0.866, p = 0.560, η^2^ = 0.010].

The concentrations of O_2_Hb, HHb, and tHb were compared before, at the end of exercise, and 10-15 minutes post-exercise (see Fig. 3, top and middle panels). The repeated sprint performance caused a significant reduction in cerebral O_2_Hb concentration (one sample *t*-test, combining GLU and PLA: t(8) = - 2.644; p = 0.029, n = 9) that was comparable between the two trials [GLU: −1.16 ± 4.13 μM; PLA: −3.11 ± 2.87 μM; F_(1, 7)_ = 2.453, p = 0.161, η^2^ = 0.096, n = 9]. On the other hand, there was a steady increase in cerebral HHb concentration during the repeated sprints (one sample *t*-test, combining GLU and PLA: t(8) = 3.248, p = 0.011, n = 9), with no effect of glucose ingestion on this response [GLU: 3.80 ± 5.95 μM; PLA: 6.44 ± 3.82 μM; F_(1, 7)_ = 1.541, p = 0.254, η^2^ = 0.042, n = 9]. Due to the increase in HHb concentration, tHb also increased after the sprint set (one sample *t*-test, combining GLU and PLA: t(8) = 3.119, p = 0.014, n = 9) regardless of glucose ingestion [GLU: 3.98 ± 5.60 μM; PLA: 5.75 ± 3.86 μM; F_(1, 7)_ = 0.670, p = 0.440, η^2^ = 0.014, n = 9].

Following the exercise, there was an increase in cerebral O_2_Hb back to baseline levels, particularly in the GLU trial [GLU: 2.54 ± 1.80 μM; PLA: 0.95 ± 2.32 μM; F_(1, 7)_ = 11.263, p = 0.012, η^2^ = 0.140, n = 9]. As a result, post-exercise O_2_Hb was higher than pre-exercise levels in the GLU trial (one sample *t*-test, t(8) = 8.099, p < 0.000, n = 9) but not in the PLA trial (one sample *t*-test, t(8) = 2.018, p = 0.08, n = 8). At 10-15 min post-exercise, HHb remained elevated in both trials [GLU: 3.42 ± 4.02 μM; PLA: 3.55 ± 3.15 μM; F_(1, 7)_ = 0.023, p = 0.884, η^2^ < 0.000, n = 9] compared to pre-exercise values (one sample *t*-test, combining GLU and PLA: t(8) = 4.557, p = 0.001, n = 9). Following changes in HHb and O_2_Hb, tHb concentration also increased after the repeated sprint exercise in both trials [Glu: 3.29 ± 4.07 μM; Pla: 3.12 ± 3.31 μM; F_(1, 7)_ = 0.050, p = 0.829, η^2^ < 0.000, n = 9], and at 10-15 min post-exercise, it was higher than at pre-exercise (one sample *t*-test, combining GLU and PLA: t(8) = 3.882, p = 0.005, n = 9).

Cardiac output and ΔO_2_Hb, ΔHHb and ΔtHb relationship

Figure 4 illustrates the relationship between cardiac output, power output, and cerebral haemoglobin concentrations (see also Supporting information, Fig. S5). During the sprints, positive correlations were found between cardiac output and the changes in O_2_Hb and ΔtHb in both GLU (cardiac output: ΔO_2_Hb, R = 0.85, p = 0.0019; cardiac output: ΔtHb, R = 0.86, p = 0.0013) and PLA trials (cardiac output: ΔO_2_Hb, R = 0.87, p = 0.0012; cardiac output: ΔtHb, R = 0.85, p = 0.0016). No significant correlation was observed between cardiac output and ΔHHb during the repeated sprints, nor between cardiac output and ΔO_2_Hb, ΔHHb and ΔtHb during the time interval between the sprints. Conversely, in the GLU trial, power output was negatively correlated with ΔO_2_Hb values during the time interval between the sprints.

**Figure 4:**
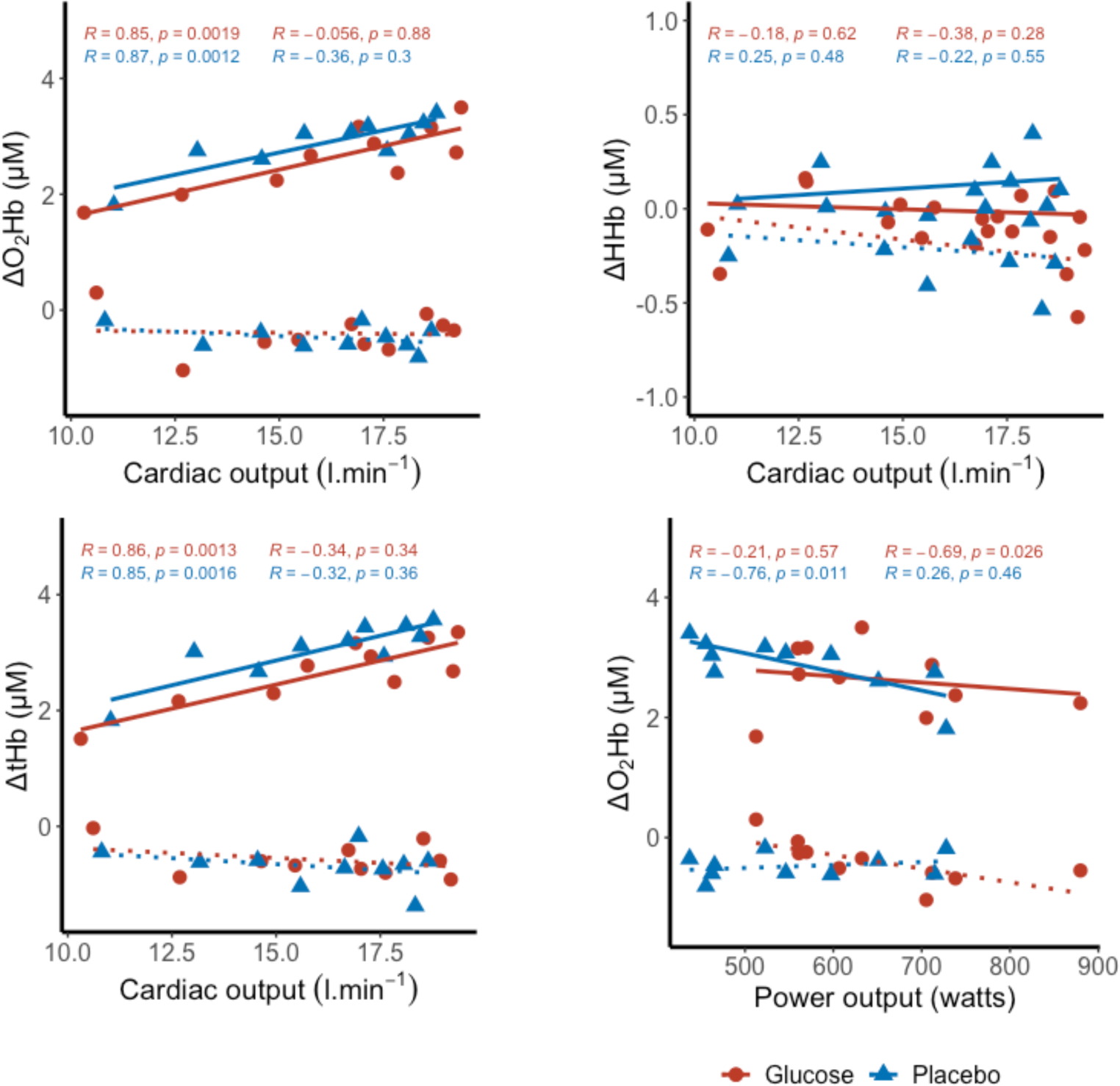
Scatter plots illustrating the relationship between ΔO_2_Hb (upper left graph), ΔHHb (upper right graph), and ΔtHb (lower left graph) with cardiac output, as well as between ΔO_2_Hb and power output (lower right graph), during the sprints (solid lines) and the time interval between sprints (dotted lines) in GLU (red lines and red circles) and PLA (blue lines and blue triangles) trials. The Pearson’s R coefficients and respective p-values are presented at the top of the graphs for values during the sprint (left column) and the time interval between the sprints (right column) for GLU (red font) and PLA (blue font) trials. During the sprint, a significant correlation was found between cardiac output and ΔO_2_Hb, as well as ΔtHb. No significant correlation was found between cardiac output and ΔHHb, and between cardiac output and ΔO_2_Hb or ΔtHb during the time interval between sprints. In the PLA trial, mean power output exhibited a negative correlation with ΔO_2_Hb during the sprints, while in the GLU trial, mean power output showed a negative correlation with ΔO_2_Hb during the time interval between sprints.

## Discussion

In this study, our aim was to outline the time profile of cerebral O_2_Hb, HHb and tHb responses to a series of brief, repeated sprints and investigate the correlation between this response and cardiac output. Furthermore, we sought to investigate the impact of acute hyperglycaemia on the changes in O_2_Hb, HHb, and tHb following the repeated sprint exercise. Our findings indicated that each sprint produced a rapid and significant surge in cerebral O_2_Hb concentration, and to a lesser extent HHb, leading to a transient increase in cerebral blood volume (i.e., increased tHb) and oxygenation. During the interval between the repeated sprints, O_2_Hb and HHb decreased and returned to, or below, pre-exercise levels rapidly. The variations in ΔO_2_Hb and ΔtHb followed a curvilinear trend similar to that observed for heart rate and cardiac output in response to the repeated sprint exercise.

During each of the short sprints, there was an increase in O_2_Hb, which was approximately ten times higher than the pulsatile peak increase due to the heartbeat at rest (∼0.5 μM). Regulation of cerebral blood flow during rest and exercise is closely tied to changes in arterial blood gases (particularly carbon dioxide), neural activity, blood pressure, and cardiac output ^21–23^. An increased cardiac output and cerebral perfusion pressure, caused by an elevation of arterial blood pressure, are two possible explanations for the transient increase in O_2_Hb and HHb that was observed during each short sprint ^12,24,25^. Although we did not record blood pressure, previous studies have shown that heavy-intensity dynamic (e.g., rowing exercise) and resistance (e.g., leg-press exercise) exercise result in a rapid increase in diastolic and systolic blood pressures, therefore it is reasonable to expect that mean blood pressure would increase during all-out sprint exercise ^12,25–27^. Although enhanced central drive and activation of the metaboreflex may contribute to the sprint-induced elevation of blood pressure ^28^, the immediate and transient increase in O_2_Hb and HHb during the short sprint exercise suggests a role for mechanical and haemodynamic factors associated with the powerful contraction of the leg and trunk muscles, resulting in brief Valsalva-like manoeuvres and synchronous elevation of arterial blood pressure ^24,27,29^. Indeed, the production of Valsalva-like manoeuvres is commonly observed during heavy static and dynamic resistance exercise dynamic exercise ^27,30^. However, this manoeuvre can also occur during dynamic exercise, such as during the catch phase of rowing ^24,29^, and is associated with the cyclical increase in MCAv in synchrony with the elevations of intra-abdominal pressure, central venous pressure, and catch force ^24^. Similarly, the onset of bilateral static leg extension simultaneously raises mean arterial blood pressure, MCAv, and cerebral O_2_Hb concentration with or without a superimposed Valsalva strain ^30^.

Previous studies have shown a direct influence of cardiac output on cerebral blood flow and cerebral oxygenation ^23,31^. In this study, we found that cardiac output had a moderate and positive correlation with changes in O_2_Hb and tHb during the sprints. This suggests that cardiac output plays a role in the rise of O_2_Hb recorded during the sprints. However, the relationship between cardiac output and cerebral blood flow is still unclear and in the present study O_2_Hb concentration declined during rest between the sprint bouts despite the increased cardiac output. While previous research has reported a direct relationship between cardiac output and MCAv during rest and moderate intensity cycling exercise (Ogoh et al., 2005), changes in MCAv during static leg exercise seem to develop independently of changes in cardiac output ^30^. Regarding our observations, we cannot refute the possibility that with increasing number of repeated cycling sprints there was an enhanced blood pressure response resulting from heightened exercise pressor reflex, central drive, and strongest Valsalva manoeuvre, thus increasing the elevation in O_2_Hb concentration across the sprints set, regardless of changes in cardiac output ^28,32^.

Despite its relationship with the changes in cerebral O_2_Hb and tHb concentration during sprints, cardiac output seems to have little influence on cerebral haemoglobin concentration levels between sprints. Following each sprint, cerebral oxygenation levels rapidly returned to baseline, while cardiac output remained elevated, indicating that a high cardiac output alone is insufficient to maintain elevated cerebral oxygenation levels. The rapid decrease in cerebral oxygenation levels with the end of each sprint could have been caused by a sudden decrease in mean arterial pressure upon cessation of the exercise ^26^. Curtelin et al. ^12^ reported a drop in mean blood pressure immediately after the end of a Wingate test, which was approximately three times larger than its increase during the effort. This decrease in mean blood pressure was accompanied by a significant decline in MCAv, which lasted for approximately 40 s before returning to pre-exercise levels ^12^. However, a decrease in CBF at the end of exercise might occur independently from changes in arterial blood pressure. For instance, Pott et al. ^24^ reported a continuous decrease in MCAv after each rowing catch movement despite stable mean blood pressure, which the authors suggest could have been caused by delayed response of dynamic cerebral autoregulation. A similar event could have taken place during the repeated sprints, given the match between the sprints’ duration and the phase delay of cerebral autoregulation ^14^.

The findings of this study showed that the repeated sprints exercise led to a progressive decrease in cerebral O_2_Hb and an increase in HHb, with considerable interindividual variation (Fig 3). The rise in HHb outweighed the decrease in O_2_Hb, resulting in an increase in cerebral tHb at the end of the repeated sprints exercise. These changes in cerebral oxygenation partially agree with MCAv data recorded during the Wingate test, revealing decreased CBF and cerebral oxygenation levels caused by augmented cerebral vascular resistance at later sprint stages ^12^. A similar response is observed during heavy resistance exercise, with cerebrovascular resistance increasing during exercise but lowering rapidly once the muscular activity stops ^26^. Thus, the data obtained from this study extends these observations, showing similarly restrained cerebral blood flow and cerebral oxygen delivery during short cycling sprints, even with moderate perceived exertion levels.

Dynamic cerebral autoregulation is a primary mechanism for regulating cerebrovascular resistance. While the effect of exercise on this mechanism is controversial, with studies indicating decreased efficacy of dynamic autoregulation during exhaustive aerobic exercise ^33^, other studies report unchanged or even enhanced cerebral autoregulation during aerobic exercise ^34^. The protracted CBF recovery from hypotension after heavy aerobic exercise ^35^ and sprinting ^12^ suggests a diminished cerebral autoregulation efficacy, possibly combined with a reduced gain of the arterial baroreflex ^35^, maybe to compensate previous mismatch between oxygen delivery and demand. While the present study cannot address the impact of short sprint performance on dynamic autoregulation, the rapid decrease of O_2_Hb at the end of the sprints when cardiac output was elevated suggests normal cerebral autoregulation during this type of exercise. Future studies may test the effect of different sprint/recovery cycle durations on cerebral haemoglobin responses. Importantly, a decline in O_2_Hb during repeated sprints contrasts with the increase in cerebral oxygenation during lower body resistance exercise ^36^. Since decreased brain oxygen saturation is observed in other sprint studies ^12^, it may be concluded that the extreme intensity of sprint exercise and the recruitment of a large muscle mass reduce cerebral blood flow and lower cerebral oxygenation. In addition, the cessation of the repeated sprints exercise was accompanied by a cerebral reactive hyperaemic response, with a steep increase in O_2_Hb within minutes following exercise (Fig. 3). This strengthens the claim that cerebral oxygen delivery is reduced during repeated sprint exercise. Finally, the absence of significant increases in both O_2_Hb and tHb during the sprint set suggests that repeated sprint performance does not cause a large increase in cerebral blood flow that could endangered the brain.

In this experiment, we investigated the effect of glucose ingestion and acute hyperglycaemia on the response of cerebral haemoglobin to sprint exercise. Although glucose is the primary energy source for the brain, hyperglycaemia has been shown to decrease the cerebral metabolic rate of oxygen ^16^ and brain connectivity ^37^ in healthy young participants at rest. Furthermore, acute and chronic hyperglycaemia decreases cerebral blood flow ^38^. In our study, we found that glucose ingestion did not affect O_2_Hb and HHb responses to the repeated sprints exercise, despite improving performance. However, glucose ingestion significantly enhanced reactive hyperaemia, resulting in faster and higher levels of O_2_Hb concentration during the post-exercise period. The relationship between post-exercise hyperaemia and any baseline changes in CBF or cerebral blood volume caused by glucose ingestion requires further investigation.

## Study limitations

There are several limitations in this study that should be acknowledged. One significant limitation is the unknown effect of extracerebral haemodynamics (scalp and skull) on cerebral oximetry measurements. Since the emitted light from NIRS travels through extracerebral tissues and the cerebral cortex, changes in blood volume and flow in superficial tissues can affect the NIRS measurements. The design of the NIRS probe used in this study, with rigid circular optodes holders firmly secured to apply pressure on the forehead skin underneath the optodes, helped to minimise changes in skin blood flow and their impact on NIRS recordings. Studies have shown that facial skin blood flow and vascular conductance increase with dynamic exercise intensity without plateauing ^39^, while ocular circulation is similar to cerebral circulation, decreasing from elevated values to baseline in response to exhaustive dynamic exercise ^40^. Based on these findings, it can be inferred that the oximetry data obtained during the repeated sprints exercise primarily reflect the changes in cerebral circulation. Future studies may assess the contribution of forehead skin to changes in haemoglobin concentration in the frontal region of the cerebral cortex during repeated sprints.

Another limitation of this study is the absence of measurements of P_ET_CO_2_ and blood pressure, which could have provided a more comprehensive understanding of the physiological changes occurring during the repeated sprints exercise.

## Perspective

We provide evidence that the performance of short cycling sprints produces a significant impact on cerebral oxygenation. We recorded a temporary increase in O_2_Hb, HHb and tHb concentrations during the sprints, with the magnitude of these changes growing with the number of sprint repetitions. In addition, we found a positive correlation between changes in O_2_Hb and cardiac output, which progressively increased throughout the repeated sprint set and remained elevated during the rest periods between sprint repetitions. However, O_2_Hb rapidly returned to baseline or below during these rest periods. Contrary to the sprint response, we observed a decrease in O_2_Hb concentration throughout the repeated sprint set. Conversely, HHb increased, suggesting inadequate brain oxygen delivery and insufficient cerebral oxygenation oxygen delivery during the all-out repeated cycling. Although not affecting haemoglobin concentration responses, glucose ingestion did improve performance and was associated with a faster recovery of O_2_Hb levels during the post-exercise period. In conclusion, our findings suggest that repeated spring exercise elicits a transient and modest increase in cerebral O_2_Hb and tHb during the physical effort itself. However, there is an overall decrease in cerebral oxygenation, leading to a robust post-exercise cerebral reactive hyperaemia in certain individuals. Importantly, our data does not indicate a marked elevation in O_2_Hb or cerebral blood flow during short all-out cycling sprints in young, non-trained individuals, thereby reducing the likelihood of cerebral hyperperfusion during this type of exercise.

## Author contributions

P.A.S. and H.M. conceptualised the study, conducted data collection, analysis, and interpretation, and drafted the original manuscript. W.Z., W.L. and F.R. contributed to conducting the experiments, data collection, and analysis. X.Z. assisted in the acquisition and analysis of NIRS data. Z.Y. and Z.K. contributed to the study conception and provided guidance in manuscript writing and revision.

## Acknowledgements

The authors thank the participants involved in this study.

