## Supplemental material for "Cerebral oxygenation and cardiac output responses during short repeated-sprints exercise and modulatory effect of glucose ingestion"

### Distribution analysis of brain oximetry data

For the statistical analysis of brain oximetry results, a log transformation was applied to the values. This decision to perform this transformation was based on the results of the Shapiro-Wilk test, which indicated non-normal distribution for several levels of the independent variables (i.e., trial, NIRS channel, and repeated sprint). To address impact of log transformation, boxplots were generated to visualize the original and log-transformed median $\Delta O_2Hb$, $\Delta HHb$, and $\Delta tHb$ values (see Figs. S1-S3). These boxplots provide a graphical representation of the data distribution and highlight any potential differences resulting from the log transformation.

*Oxyhaemoglobin*

Figure S1 provides boxplots illustrating the distribution of median $\Delta O_2Hb$ values (A) and their corresponding natural log-transformed values (B) across each channel and repeated sprint. The analysis revealed the presence of outliers within the dataset. Out of a total of 1600 $\Delta O_2Hb$ measures, 111 were identified as outliers, with 30 of them classified as extreme outliers. Upon applying the logarithmic transformation to the data (Fig. S1, B), the variability in median $\Delta O_2Hb$ values decreased as expected. However, a considerable number of outliers (80 out 1600 observations) remained, with the number of extreme outliers reduced to 13.

A chi-squared test was performed to compare the proportion of outliers between non-transformed and log-transformed data, considering the two trials and eight channels. The results indicated different proportions of outliers depending on the sprint repetition and individuals for both untransformed and log-transformed $\Delta O_2Hb$ values (Table S1). Notably, a visual examination of the boxplots revealed an apparent increase in the number of outliers with an increasing number of sprint repetitions. Additionally, the proportion of outlying $\Delta O_2Hb$ measures differed among participants, with some participants contributing to a substantial proportion of the total outliers, while others had no outlying measurements.

*Deoxyhaemoglobin*

Figure S2 displays the boxplots depicting the distribution of median ΔHHb values (A) and their corresponding natural log-transformed values (B) across each channel and repeated sprint. Similar to $\Delta O_2Hb$, a significant number of outliers were observed for ΔHHb values. In total, 147 outliers were identified for both non-transformed and log-transformed ΔHHb measures. Among these outliers, 54 and 52 outliers were classified as extreme for non-transformed and log-transformed ΔHHb data, respectively. The proportion of outliers remained consistent across the two trials, channels, and repeated sprints (Table S1). However, the distribution of outliers varied among participants, suggesting individual variations in ΔHHb measurements.

*Total haemoglobin*

Figure S3 presents the boxplots representing the median tHb values (A) and their log-transformed counterparts (B). The analysis revealed a substantial number of outlying tHb values, with a total of 120 outliers, including 37 extreme outliers. The tHb outliers were evenly distributed across the PLA and GLU trials and NIRS channels; however, their distribution varied with the different repeated sprints (Table S1). Examining Fig. S3 reveals that the number of tHb outliers increased as the number of sprint repetitions progressed, with the highest number of outliers observed during the final two repeated sprints, with 19 outliers in each. Log-transforming the tHb data resulted in a reduction of outliers, with a total of 98 outliers, and 20 extreme outliers. The proportion of outliers after log-transformation was consistent across trials, NIRS channels, and repeated sprints (Table S1).

Table S1: Results of $\chi^2$ (df) tests for the proportion of outliers for $\Delta O_2Hb$, $\Delta HHb$, and $\Delta tHb$ according to the trial, NIRS channel, repeated sprint, and participant.

| | | Non transformed values | Log-transformed values |
|---|---|---|---|
| $\Delta O_2Hb$ | Trial | $\chi^2$ (1) = 0.081, p = 0.776 | $\chi^2$ (1) = 0.111, p = 0.739 |
| | Channel | $\chi^2$ (7) = 3.52, p = 0.833 | $\chi^2$ (7) = 5.815, p = 0.562 |
| | Sprint | **$\chi^2$ (9) = 32.87, p < 0.000** | **$\chi^2$ (9) = 24.06, p < 0.004** |
| | Participants | **$\chi^2$ (9) = 263.32, p < 0.000** | **$\chi^2$ (9) = 93.198, p < 0.000** |
| $\Delta HHb$ | Trial | $\chi^2$ (1) = 3, p = 0.083 | $\chi^2$ (1) = 1.97, p = 0.161 |
| | Channel | $\chi^2$ (7) = 3.37, p = 0.849 | $\chi^2$ (7) = 4.02, p = 0.777 |
| | Sprint | $\chi^2$ (9) = 14.29, p = 0.112 | $\chi^2$ (9) = 14.29, p = 0.112 |
| | Participants | **$\chi^2$ (9) = 142.32, p < 0.000** | **$\chi^2$ (9) = 144.36, p < 0.000** |
| $\Delta tHb$ | Trial | $\chi^2$ (1) = 0.53, p = 0.465 | $\chi^2$ (1) = 0.04, p = 0.840 |
| | Channel | $\chi^2$ (7) = 2.67, p = 0.944 | $\chi^2$ (7) = 7.31, p = 0.398 |
| | Sprint | **$\chi^2$ (9) = 24.5, p = 0.004** | $\chi^2$ (9) = 14.653, p = 0.101 |
| | Participants | **$\chi^2$ (9) = 188.67, p < 0.000** | **$\chi^2$ (9) = 70.367, p < 0.000** |

Figure S1

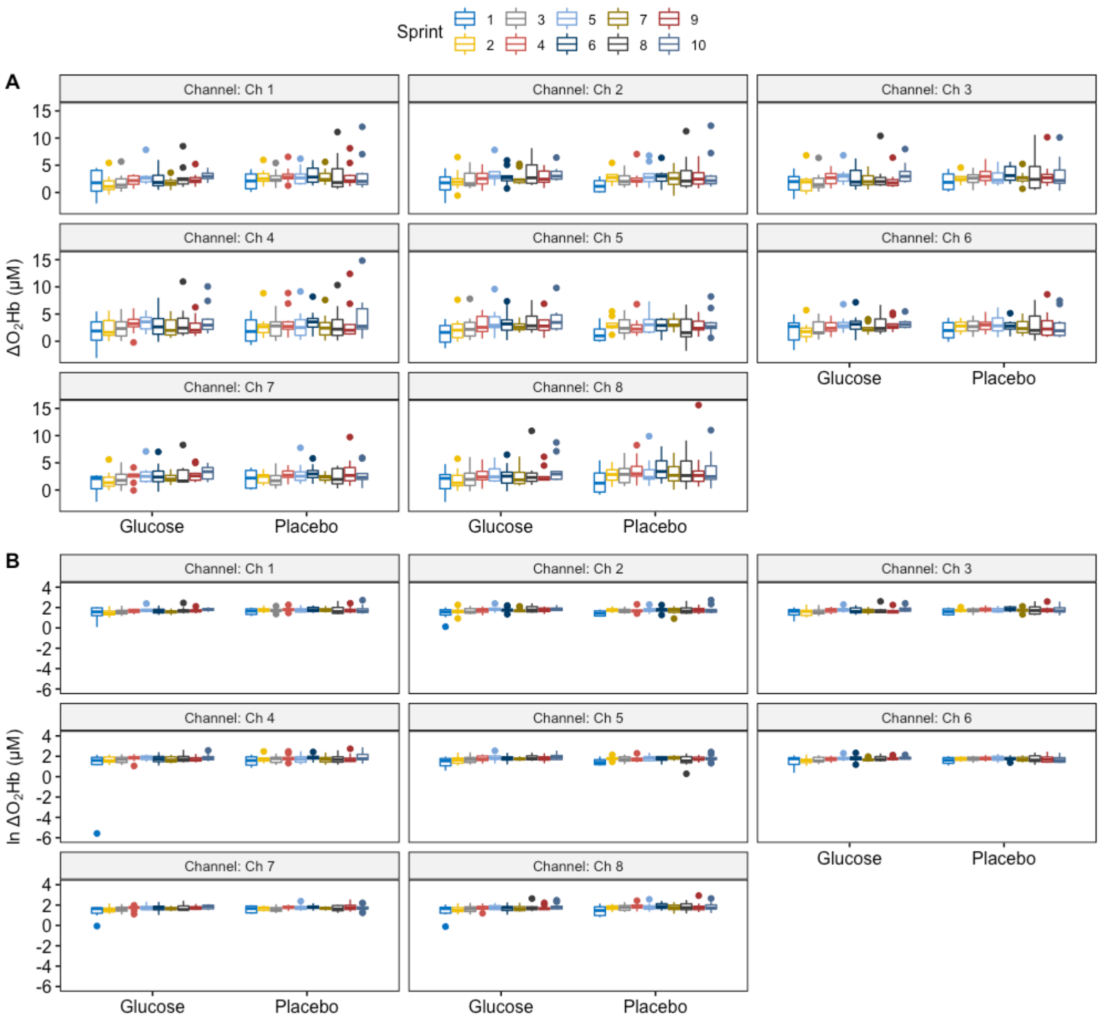


Figure S1: Boxplots for non-transformed (A) and natural log-transformed (B) median $\Delta O_2Hb$ data for each channel and repeated sprint.

Figure S2

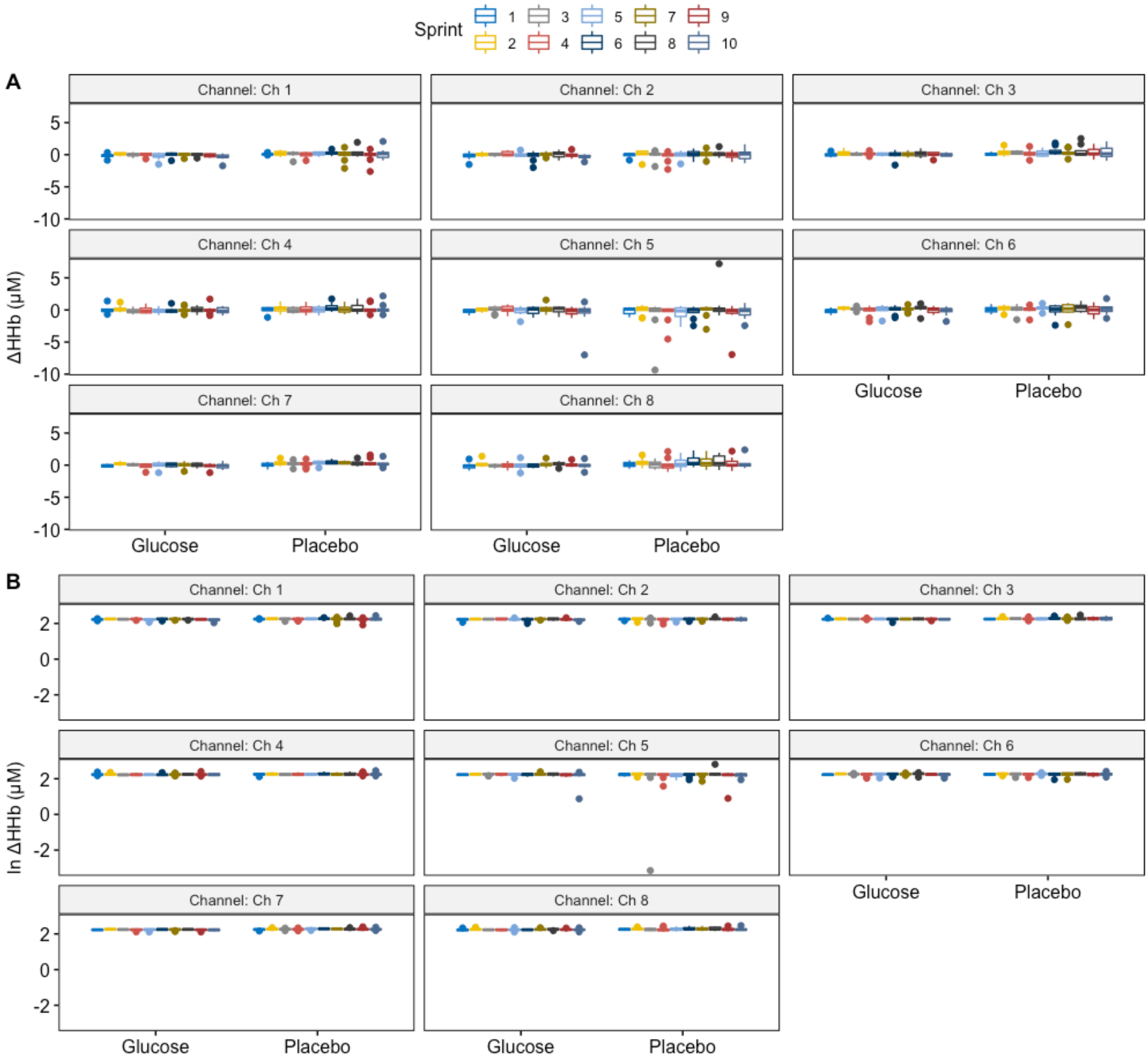


Figure S2: Boxplots for non-transformed (A) and natural log-transformed (B) median ΔHHb data for each channel and repeated sprint.

Figure S3

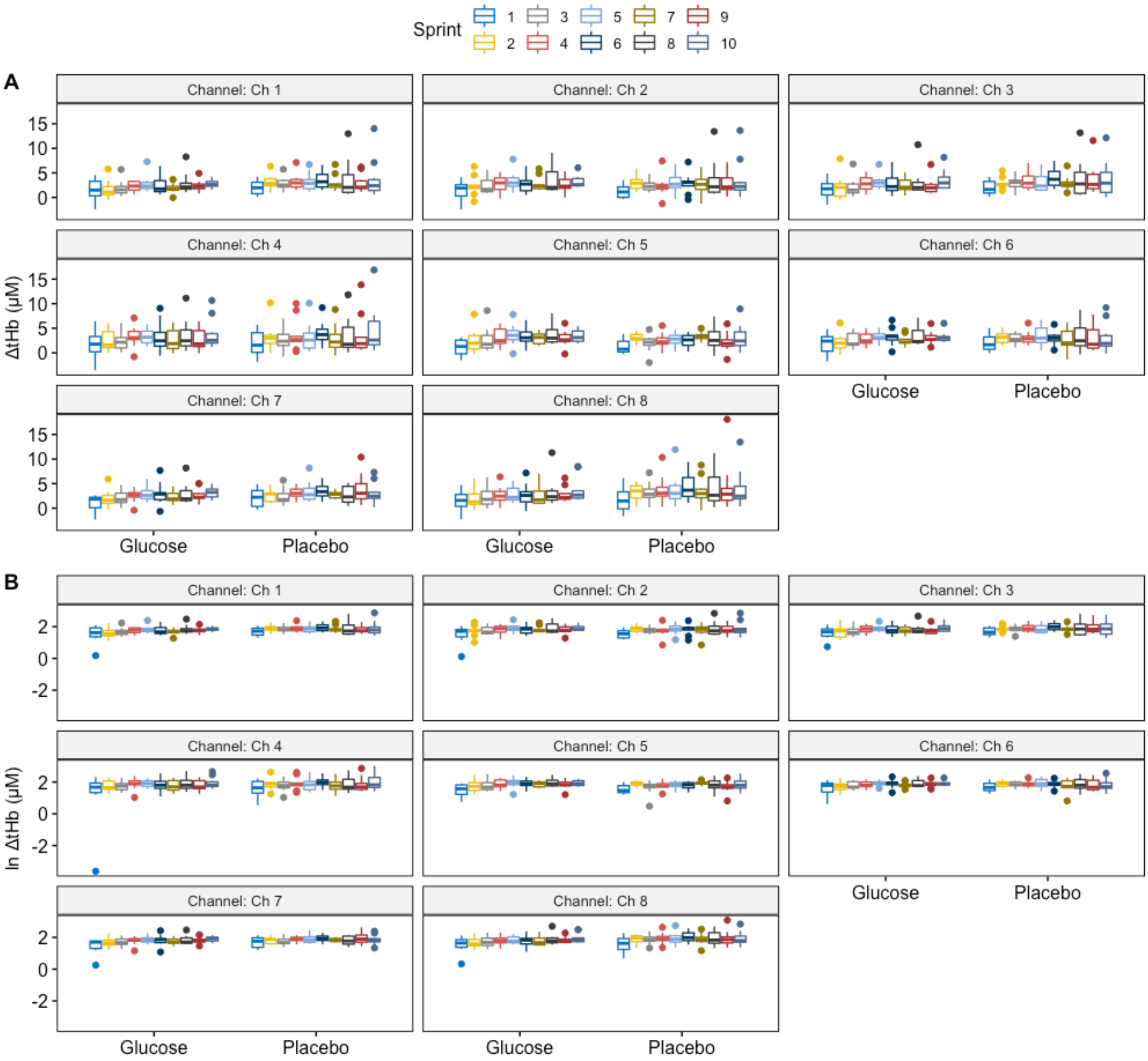


Figure S3: Boxplots for non-transformed (A) and natural log-transformed (B) median ΔtHb data for each channel and repeated sprint.

Figure S4

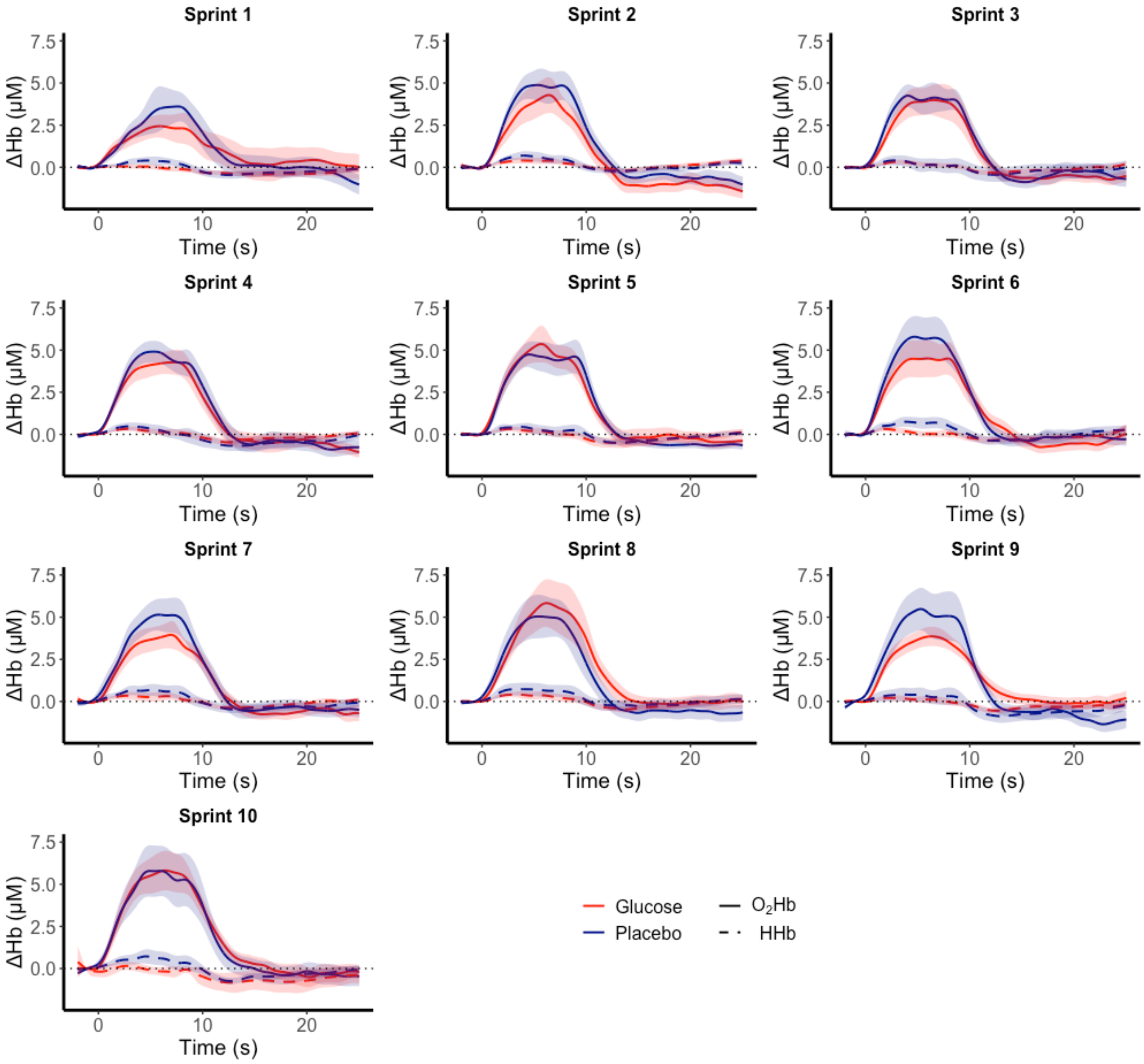


Figure S4: Changes in $O_2Hb$ (solid lines) and HHb (dashed lines) concentrations during repeated sprints for GLU (red lines) and PLA (blue lines) trials. Time zero is the start of each sprint. Notably, a transient $O_2Hb$ and HHb increase is observed in all repeated sprints. At time ~15 s, both $O_2Hb$ and HHb concentrations return to baseline or slightly below. Also shown is the smaller increase in $O_2Hb$ during the first sprint, as well as the progressive increase in $O_2Hb$ responses with each subsequent sprint repetition.

Figure S5

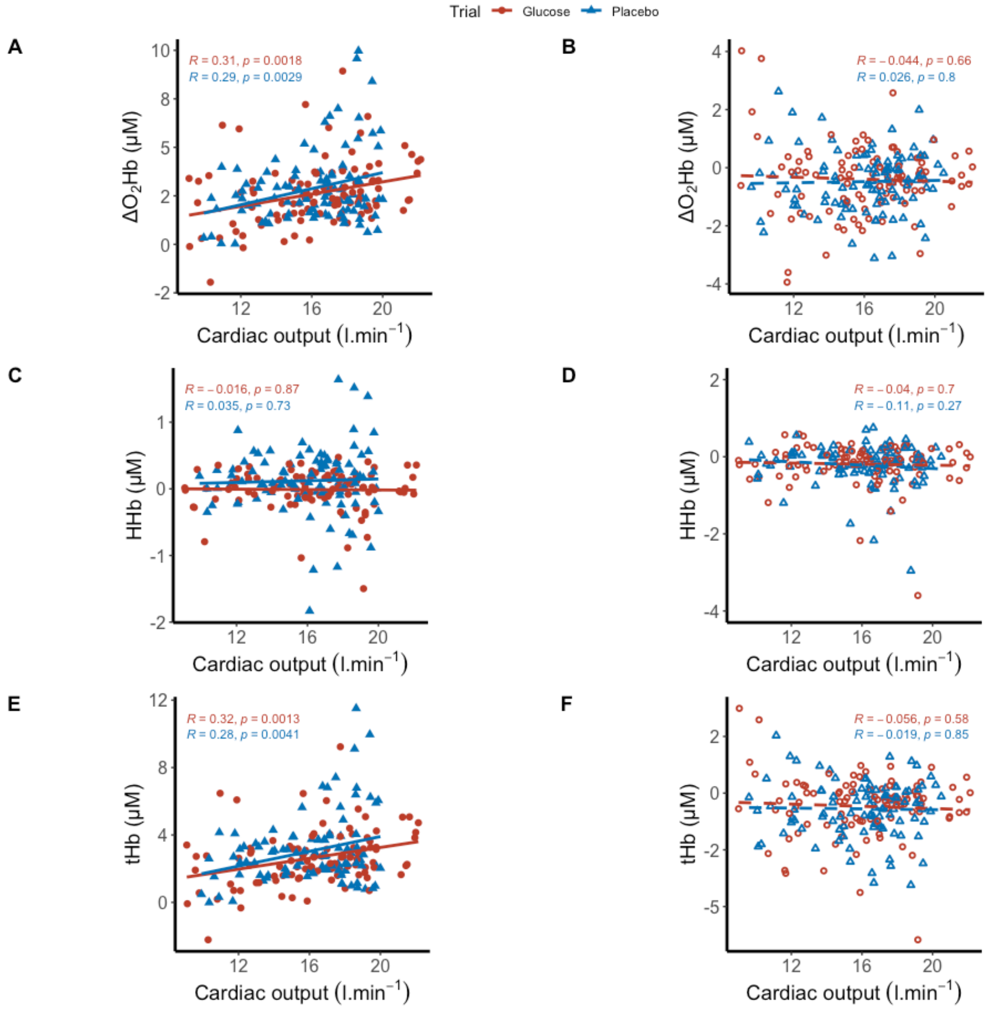


Figure S5: Scatter plots showing the relationship between cardiac output and median $\Delta O_2Hb$ (A, B), ΔHHb (C, D), and ΔtHb (E, F) during the repeated sprints (A, C, E) and during the time interval between sprints (B, D, F) for the GLU and PLA trials. Each data point represents the average of the eight NIRS channels for each participant and repeated sprint in the two trials (200 data points). The Pearson's R coefficients and respective p-values are indicated above each graph using the same colour scheme as the data points and lines. Strong positive correlations were observed between median $\Delta O_2Hb$ and ΔtHb during the sprints and cardiac output (A, E). No significant relationship was found for ΔHHb and cardiac output during the sprints (C), nor between cardiac output and any of the haemoglobin concentration changes during the time interval between sprints (B, D, F).
